# Low-coverage sequencing in a deep intercross of the Virginia body weight lines provides insight to the polygenic genetic architecture of growth: novel loci revealed by increased power and improved genome-coverage

**DOI:** 10.1101/2021.07.19.451141

**Authors:** T. Rönneburg, Y. Zan, C.F. Honaker, P.B. Siegel, Ö. Carlborg

**Affiliations:** Department of Medical Biochemistry and Microbiology, Uppsala University, Uppsala, Sweden; Department of Animal and Poultry Sciences, Virginia Polytechnic Institute and State University, Blacksburg VA, USA

## Abstract

Genetic dissection of highly polygenic traits is a challenge, in part due to the power necessary to confidently identify loci with minor effects. Experimental crosses are valuable resources for mapping such traits. Traditionally, genome-wide analyses of experimental crosses have targeted major loci using data from a single generation, often the F_2_, with additional, later generation individuals being generated for replication and fine-mapping. Here, we aim to confidently identify minor-effect loci contributing to the highly polygenic basis of the long-term, divergent bi-directional selection responses for 56-day body weight in the Virginia chicken lines. To achieve this, a powerful strategy was developed to make use of data from all generations (F_2_-F_18_) of an advanced intercross line, developed by crossing the low and high selected lines after 40 generations of selection. A cost-efficient low-coverage sequencing based approach was used to obtain high-confidence genotypes in 1Mb bins across 99.3% of the chicken genome for >3,300 intercross individuals. In total, 12 genome-wide significant and 10 additional suggestive QTL for 56-day body weight were mapped, with only two of these QTL reaching genome-wide, and one suggestive, significance in analyses of the F_2_ generation. Five of the significant, and four of the suggestive, QTL were among the 20 loci reaching a 20% FDR-threshold in previous analyses of data from generation F_15_. The novel, minor-effect QTL mapped here were generally mapped due to an overall increase in power by integrating data across generations, with minor contributions from increased genome-coverage and improved marker information content. Significant and suggestive QTL now explain >60% of the difference between the parental lines, three times more than the previously reported significant QTL. Making integrated use of all available samples from multiple generations in experimental crosses is now economically feasible using the low-cost, sequencing-based genotyping strategies outlined here. Our empirical results illustrate the value of this strategy for mapping novel minor-effect loci contributing to complex traits to provide a more confident, comprehensive view of the individual loci that form the genetic basis of the highly polygenic, long-term selection responses for 56-day body weight in the Virginia chicken lines.

## Introduction

Quantitative traits remain difficult to analyse and break down into their component loci (Flint and Mott 2001). Effect sizes of individual loci often explain a very small fraction of the phenotypic variance (Boyle, Li, and Pritchard 2017 and references within) - often much smaller than environmental effects - and are regularly dependent on the genetic background (Pettersson et al. 2011; Mackay 2014; Forsberg et al. 2017; Zan and Carlborg 2020). Experimental populations are valuable resources for studying quantitative traits, and by reducing confounding factors such as environmental noise, they have provided a clearer view on the genetic architecture of a wide range of complex traits (Flint and Mott 2001; Andersson 2001; Andersson and Georges 2004). Examples include shank-length in mice (Castro et al. 2019), longevity in Drosophila melanogaster (Curtsinger and Khazaeli 2002), and oil content in corn (Hopkins 1899; Dudley 2007).

Although QTL (Quantitative trait loci) studies in experimental crosses have high power, resolution is limited due to the extensive LD (Linkage disequilibrium) introduced by the crossing design. Historically, sparse or very sparse marker maps have therefore been used, resulting in regions of the genome with less coverage and, in some cases, missing data on small chromosomes and/or chromosomal ends (Mackay 2001). Similarly, it is not uncommon to have regions lacking markers informative for line origin when using experimental crosses between outbred founders (Andersson 2001). To increase resolution and facilitate fine-mapping of detected QTL, follow-up studies in additional crosses have been performed. Generally, these have excluded regions outside of previously observed QTL, therefore leaving much of the genome without further study. As a result, these studies are underpowered or lack the resolution to make inferences on the genetic architecture of the studied traits beyond a few large-effect loci (Flint and Mott 2001).

More complete dissection of highly polygenic complex traits requires large and powerful studies. In natural populations such as humans, hundreds of thousands of individuals have been used to study highly polygenic model traits such as height (Lango Allen et al. 2010; Yang et al. 2010; Wood et al. 2014). In experimental populations, smaller populations are required to detect even minor-effect loci due to the higher power achieved from, for example, segregation of alleles at intermediate frequencies and greater control over environmental influences. New genotyping and imputation approaches based on low-coverage whole genome sequencing (WGS; Altshuler et al. 2000; Andolfatto et al. 2011; Zhang et al. 2015; Pértille et al. 2016; Whalen et al. 2018; Zan et al. 2019), provide opportunities to reanalyse existing individuals generated for different purposes to perform integrated analyses, thus enabling greater insights into the contribution of loci with minor effects on the genetic basis of complex traits in existing experimental populations. The increased genome-wide marker coverage provided by WGS-based genotyping technologies also provides an extended coverage of regions outside of the current consensus linkage maps in species such as the chicken, where, for example, the major focus has been on the large chromosomes, leaving the microchromosomes largely unexplored (Groenen et al. 2000; 2009).

The Virginia body weight lines of chickens were developed by long-term, bi-directional selection for a single trait – body weight at 56 days of age – resulting in a nine-fold difference between the low (LWS) and high (HWS) lines after 40 generations of selection (Dunnington and Siegel 1996; Dunnington et al. 2013; Márquez, Siegel, and Lewis 2010). Genome-wide comparisons showed that the footprint of selection between the LWS and HWS cover hundreds of loci across the genome (Johansson et al. 2010; Lillie et al. 2018; Lillie et al. 2019). Efforts to identify which of these loci contribute to the observed responses include a series of experiments utilizing an intercross developed from individuals of generation S_41_ (n_HWS_=29, n_LWS_=30). Efforts include genome-wide mapping (Jacobsson et al. 2005; Carlborg et al. 2006; Wahlberg et al. 2009), as well as replication and fine-mapping studies (M. Pettersson et al. 2011; Besnier et al. 2011; Sheng et al. 2015; M. E. Pettersson et al. 2013; Brandt et al. 2017; Zan et al. 2017) on different generations in this population. Although these studies agree that the long-term responses are primarily from selection on a highly polygenic genetic architecture where most loci have small effects, the statistical support for individual loci is low. This study aims to overcome this deficiency of statistical power to facilitate the mapping of contributing minor-effect loci with confidence by re-genotyping and performing an integrated analysis of >3,300 individuals from generations F_2_-F_18_ of the Virginia lines intercross, identifying and mapping new QTL, confirming earlier reported loci, and explaining more of the selection responses with individually significant loci than previously reported, thus illustrating the value of utilizing new and affordable WGS-based genotyping strategies.

## Methods

### *Deep-Stripes*: a pipeline for founder-line genotype estimation in deep intercross populations

*Stripes* (Zan et al. 2019) is a pipeline for founder-line genotype estimation using low-coverage sequencing data, extending *TIGER* (Rowan et al. 2015) for use in outbred intercross populations. In deep intercross populations from outbred founders, such as the advanced intercross line (AIL) studied here, there is a generally lower and more variable density of founder-line informative markers. Here, we have further extended the *Stripes* pipeline to deep intercross populations, including updates to enhance stability, and improve genotype calling quality in later generations.

*Deep-Stripes* updates are implementations of (a) reverting to hardcoded genotype emission thresholds in cases where the original nonlinear minimisation procedure for determining these (Rowan et al. 2015) failed due to uniform ancestry across an entire chromosome, (b) a modified nonlinear minimisation procedure improves convergence as well as defaulting to hardcoded parameters after 20 unsuccessful tries to determine the genotype emission thresholds, (c) a modified logic for comparing highly similar beta distributions to make results stable across computing platforms, and (d) automation of multiple rounds of genotype estimation for each individual (forward and reverse on each chromosome with an arbitrary number of window sizes - here 50 and 200 markers).

### Genotype quality control and filtering

*Deep-Stripes* implemented genotype estimation in both directions on the chromosome. This facilitated detection of incoherently called genotypes in low-information areas due to a delay of inferred crossovers to the end of such regions. Genotype estimation with two window-sizes was used to reduce the number of false positive crossovers in marker-dense areas resulting from the flat, per-window genotype estimation error rate.

Final genotype estimation was done by transforming the output from each of the four runs described above to a genotype matrix. These contained estimated founder line genotypes for each individual in even-sized (1 Mb) bins across the genome. These four matrices were processed as follows: In bins where no recombination event was inferred, the genotypes were coded as numerical values (1, 0, −1) corresponding to the homozygote for founder-line 1, heterozygote and homozygote for founder-line 2, respectively. Recombination breakpoints were estimated with bp resolution, but if one or more recombination events were detected, resulting in multiple genotypes being called in a 1 Mb window, the genotype was scored on a continuous scale from 1 to −1 by averaging the founder genotypes scores across the base pairs in the segment. Second, the genotype matrices from the forward and reverse runs were filtered by considering bins where estimates differed by more than one recombination event as uninformative and setting them to missing. This procedure was performed separately for the two window-sizes (50 and 200 markers). Third, the forward- and reverse-filtered genotype matrices obtained using 50 and 200 marker window-sizes were combined. This was done by using the genotype derived using 50/200 marker windows, bins with ambiguous genotypes were set to missing. They were defined as bins with genotype scores in the ranges [0.8,0.2] and [−0.8, −0.2]. Finally, all bins genotyped in less than 100 individuals were set to missing.

### Genotype estimation in the Virginia lines AIL using *deep-Stripes*

The *deep-Stripes* pipeline described above was used to call founder line origin genotypes in 1 Mb bins across the genome of the F_2_-F_18_ generations of the AIL. For this, high-coverage sequence data from the outbred founders of the population and low-coverage sequence data from the intercross individuals generated as described below was used.

### Founder-line sequencing and variant calling

All 59 high (HWS) and low (LWS) founders of the AIL (n_HWS_ = 29 and n_LWS_ = 30) were whole-genome re-sequenced to ~30X coverage (Guo et al. 2019). The obtained reads were then mapped to the newest reference genome (GalGal6a, Genome Reference Consortium 2018) using *BWA* (version 0.7.17 (Li 2013)). Variants were called and filtered using GATK (McKenna et al. 2010) according to best-practices recommendations (DePristo et al. 2011; Auwera et al. 2013) modified to accommodate for non-model organisms. The code and parameters used for this analysis are provided in the Supplementary File 1 and the associated Github repository (github.com/CarlborgGenomics/AIL-scan).

### Low-coverage sequencing of advanced intercross line individuals

All chickens from generations F_2_-F_18_ of the AIL (n_F2-F18_ = 3,327, Table S2) were sequenced to ~0.4X coverage (Zan et al. 2019). The obtained reads were mapped to the GalGal6 reference genome using *BWA* (version 0.7.17, Li 2013). Variants were next called using a pipeline implemented using *bcftools* (1.9,Li 2011), *samtools* (1.9,Li et al. 2009), *biopython* (1.70, Cock et al. 2009), and *cyvcf2* (0.10.0, Pedersen and Quinlan 2017). The code and parameters used are provided in Supplementary File 2 and the project github repository. Obtained variants were merged for each generation and filtered to only include polymorphisms present in the filtered set of founder genotypes described in the section above. Next, for each individual, only variants informative about the founder line origin (HWS/LWS) were kept and formatted for use as input to the *Stripes* genotyping pipeline (Zan et al. 2019).

### QTL mapping

QTL mapping was performed for body weight at 8 weeks of age. To correct for generational effects, 8-week body weights were standardized within each generation. This was done by subtracting the generation mean from each observation and then dividing it by the within-generation standard deviation. The genome scan was performed using the ‘scanone’ function from the package ‘qtl’ (Broman et al. 2003) in R (R Core Team 2013) using Haley-Knott regression (Knott and Haley 1992) with sex as a covariate. Significance threshold was obtained using permutations (n=10,000), resulting in a 5% genome-wide significance of LOD = 4.01. For suggestive significance, the 5% chromosome-wide significance on Chromosome 4 was taken to be consistent with previous studies (LOD = 2.86; Jacobsson et al. 2005; Wahlberg et al. 2009).

### Selection of an extended marker set using FDR approach

In order to obtain a larger, more lenient set of markers, LOD scores were transformed into p-values (Peirce et al. 2006). They were then evaluated against significance thresholds adjusted for multiple testing using the Benjamini Hochberg procedure with a false discovery rate of 10%, as implemented in *statsmodels* (Benjamini and Hochberg 1995; Seabold and Perktold 2010).

### Estimation of the genetic effect and residual variance explained by the mapped QTL

To estimate the residual variance explained by the QTL, we corrected for the sex effect using a linear model and fit either i) all significant or ii) all suggestive and significant QTL using the *fitqtl* function in *r/qtl* on the residuals. Estimates for the residual variance explained by each QTL were obtained by fitting all significant and suggestive QTL jointly and using the SSv3 drop-one-term anova in the *fitqtl* function. Estimates for the effect on body weight in grams for each QTL were determined by fitting each QTL individually with sex as a covariate. Due to the standardisation of the phenotypes, the estimates were multiplied with the population-standard deviation to obtain an estimate in grams. The sum of these estimates was then expressed as a fraction of the between-line difference.

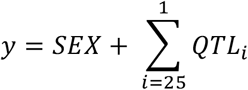

### Estimating the effect of increased genotyping information content on statistical power in QTL analyses

The information content (IC) was calculated across the genome in the set of F_2_ individuals that were common to this study and that of Wahlberg et al. (2009). The measure used was defined as (Knott et al. 1998):

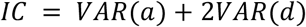

This was calculated at individual marker locations, as well as every cM, across the genome using the *a* and *d* indicator regression variables from Wahlberg et al. (2009). For the dataset from this study, the a and d indicator variables were calculated from the genotype estimates in each 1Mb bin as (Knott and Haley 1992):

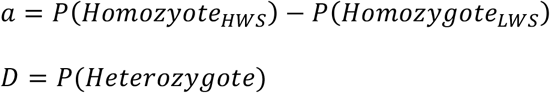

The information content was compared at the physical locations (Mb, Genome Reference Consortium 2018) of the genotyped markers in Wahlberg et al. (2009) and across all tested locations in the two studies (every cM/Mb; Wahlberg et al. 2009/this study).

## Results

### Increased coverage in the genome-wide scan via genotyping by low-coverage sequencing

After sequencing and variant calling, genotypes were estimated for n_F2-F18_ = 3,327 AIL individuals that passed quality control. The average density of informative markers across the genome was 102 markers/Mb, though since the founder lines are outbred and the individual AIL offspring descend from different founders, the number of informative SNPs varies among individuals and decreases over generations, as more ancestors contributed to the genotype of each individual. Founder-line genotypes were obtained for all of the 1,058 1Mb bins defined on the 33 largest chromosomes, identifying an average of 74 recombination breakpoints per Mb across all individuals, before filtering. Compared to the previous genome-wide scan performed in the F_2_ population by Wahlberg et al. (2009), an additional 100 Mb were covered (+27%) with markers, encompassing two small chromosomes (Chr31, Chr33), several previously uncovered scaffolds/unplaced segments and chromosome ends (Figure 1). Further, the information content at the tested locations across the genome also improved from an average of 0.77 (Wahlberg et al. 2009) to an average of 0.90 (Figure 3, panel F).

**Figure 1.**
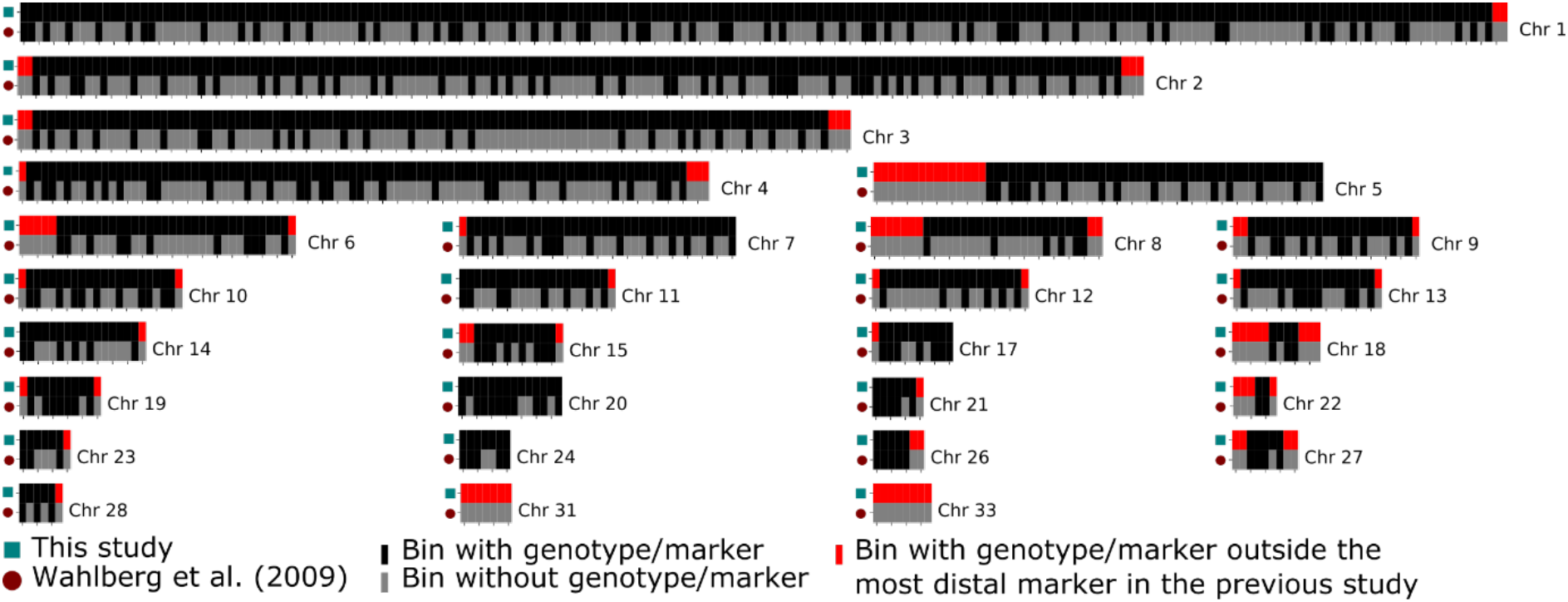
Genome coverage by the use of low-coverage sequencing data compared to an earlier F_2_ genome scan (Wahlberg et al. 2009) based on 372 SNP and microsatellite markers. Black/grey colors indicate 1 Mb bins with/without genotypes and red highlights chromosome ends with new genotypes outside the outermost markers in Wahlberg et al. (2009).

**Figure 2:**
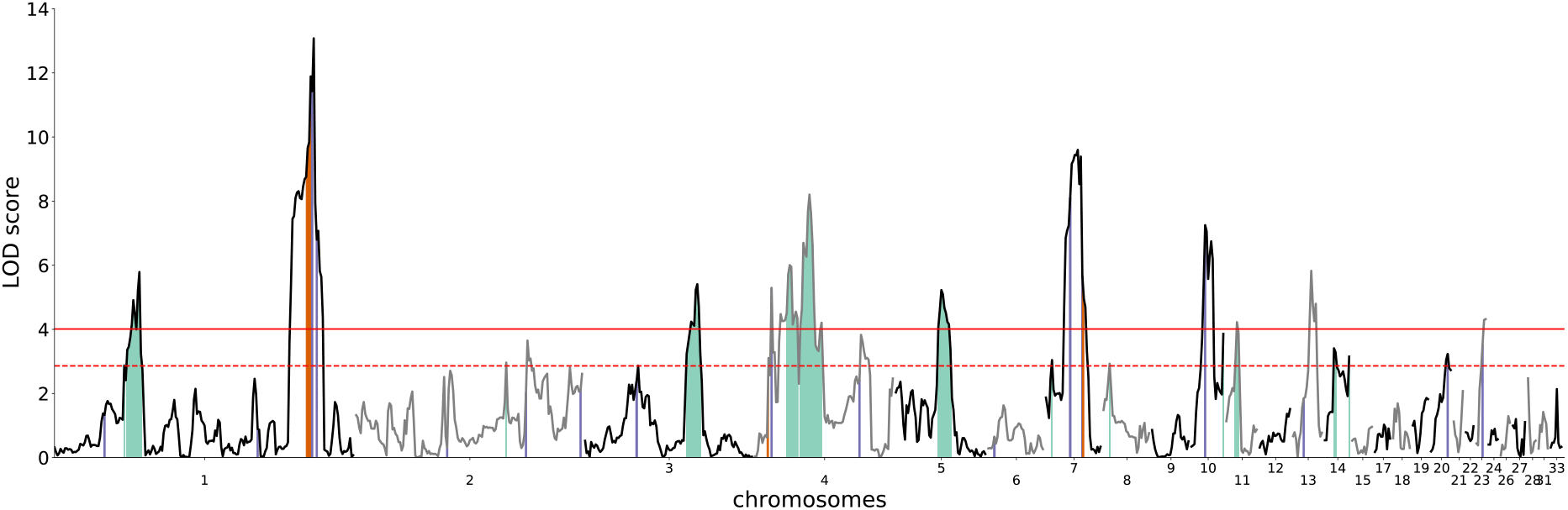
Genome-wide QTL scan for 56-day body weight in generations F_2_-F_18_ of the Virginia body weight lines AIL. The y-axis shows the statistical support for QTL as LOD scores and the x-axis the genomic location in Mb bins. The solid/dashed red horizontal lines show the genome-wide/suggestive significance thresholds, respectively. Red vertical segments show the most recent earlier reported associations in genome-wide scans in the F_2_ (Wahlberg et al. 2009), blue vertical segments indicate associations in the fine-mapping analyses in the *F*_15_ (Zan et al. 2017). Green vertical segments indicate new suggestive QTL without a previous association within 15 Mb.

**Figure 3:**
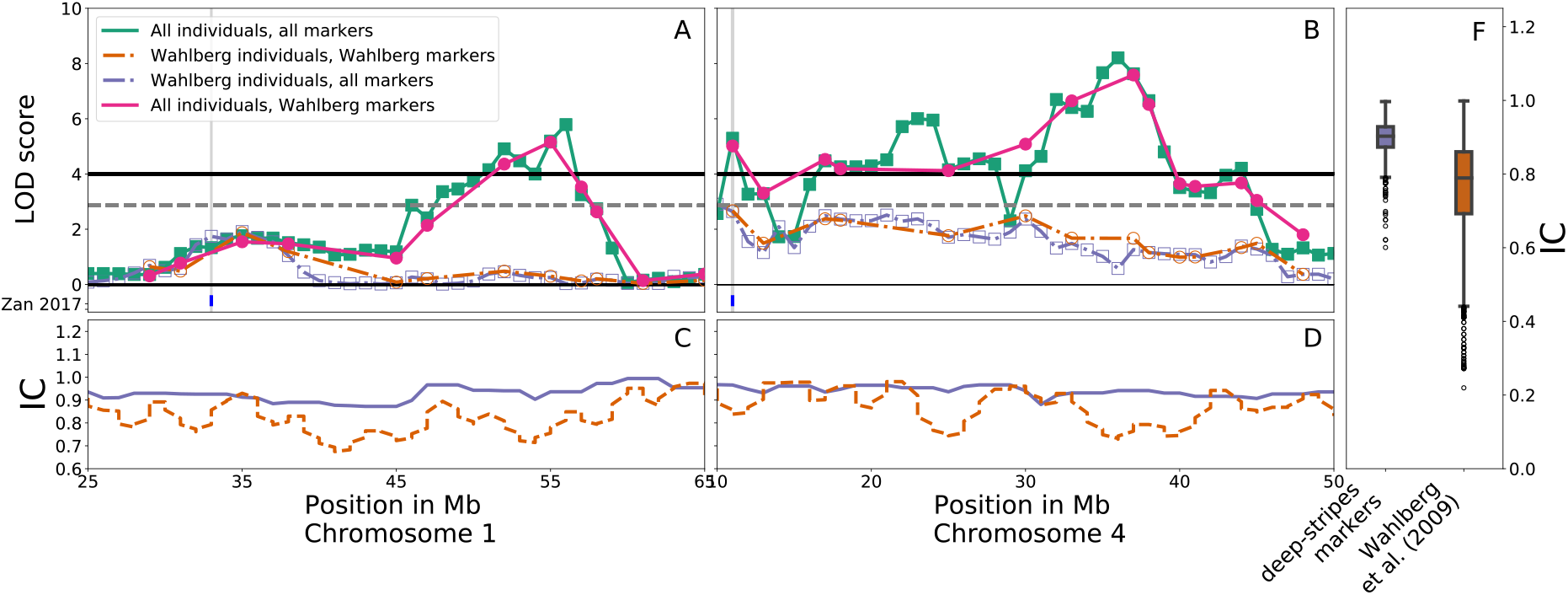
Upper left and middle panels show the LOD score across selected peak regions on chromosome 1 (25-65 Mb) and chromosome 4 (10-50 Mb), using different selections of markers and individuals. Dashed/solid lines show the QTL scans using the F_2_ individuals from Wahlberg et al. / All individuals, solid round / empty square markers show QTL scans using markers from Wahlberg et al. (2009) / low coverage data. Bottom left and middle panels display information content across the same regions on chromosome 1 and 4 for the markers used in Wahlberg et al. (orange, dashed) and the *deep-Stripes* markers (periwinkle, solid) across the individuals used by Wahlberg et al. (2009). The right panel summarises information content for the same sets of markers and individuals, but across the 30 largest chromosomes.

### More individuals and increased genome-coverage facilitate detection of new QTL

#### Comparisons to earlier mapped and fine-mapped QTL in the Virginia lines AIL

As illustrated in Figure 2, the 2+1 genome-wide significant and suggestive QTL in the most recent genome-scan of the F_2_ population (Table 1; Wahlberg et al. 2009) were detected. In addition, nine additional QTL for this trait were mapped with genome-wide significance (Figure 2; Table 1). Using the FDR-adjusted threshold for significance, a total of 42, 33, 21 QTL were identified at false discovery rate of 10%, 5% and 1%, respectively (Fig. S3, Tab. S3). This resulted in 20 QTL in addition to the QTL reaching genome or chromosome wide significance using the permutation approach.

**Table 1:** Significant QTL for 56-day body weight and overlaps with earlier reported significant and suggestive QTL in this population.

| Chromosome | Position (Mb) | a(SE) | %var | LOD | QTL | previous |
| --- | --- | --- | --- | --- | --- | --- |
| 1 | 56 | 19.6(4.0) | 0.43 | 5.79 |  |  |
| 1 | 171 | 31.6(4.1) | 1.21 | 13.08 | Growth1 | Zan 2017, Wahlberg 2009, |
| 3 | 74 | 20.3(4.2) | 0.31 | 5.41 | Growth4 |  |
| 4 | 11 | 20.7(4.4) | 0.28 | 5.29 | Growth6 | Zan 2017,Wahlberg 2009 |
| 4 | 23 | 20.4(4.0) | 0.07 | 6 | Growth7 |  |
| 4 | 36 | 25.6(4.2) | 0.45 | 8.2 | Growth7 |  |
| 5 | 30 | 20.9(4.3) | 0.31 | 5.22 | Growth8 |  |
| 7 | 21 | 25.6(4.0) | 1.26 | 9.6 | Growth9 | Zan 2017, Wahlberg 2009 |
| 10 | 9 | 22.3(3.9) | 0.27 | 7.25 |  | Zan 2017 |
| 11 | 7 | 13.2(4.0) | 0.32 | 4.22 |  |  |
| 13 | 12 | 19.6(3.9) | 0.32 | 5.83 | Growth10 |  |
| 23 | 6 | 10.9(4.3) | 0.47 | 4.32 |  | Zan 2017 |
|  |  | a=250.7 |  |  |  |  |
|  |  | 2a=501.4 |  |  |  |  |

#### Overlap with suggestive regions

Previously, fine-mapping had been performed in significant and suggestive QTL-regions and identified selective sweep regions (Besnier et al. 2011; Sheng et al. 2015; Zan et al. 2017). In contrast to the original F_2_ genome-scan (Wahlberg et al. 2009), two of the fine-mapped loci (located on Chromosome 10, Mb 9 and 23, Mb 6) located outside of the 2+1 originally significant and suggestive QTL reached genome-wide significance when analysed across the entire AIL (Figure 2, Table 1), with four more finemapping loci overlapping QTL that reach suggestive significance in this study (4/70, 2/113, 3/35, 10/11 Chr/Mb). When comparing the extended set of 42 QTL to the markers from Sheng et al. (2015) which tag a set of 99 regions under selection that were identified by Johansson et al. (2010), 41 of the regions overlapped the extended list of QTL with at least one tagging marker.

#### More individuals and increased marker density dissects a QTL on Chromosome 4 into multiple, independent associated regions

Previously, two regions on Chromosome 4 were implicated for body weight, with one of them, *Growth6*, reaching genome wide significance for 56-day body weight in the latest genome scan (Wahlberg et al. 2009). The population used here provided sufficient power to replicate the previously reported QTL and confirm the association of another region, *Growth7*, that was previously reported as associated with body weight and growth traits (Figure 3, panel B, compares F_2_ individuals from Wahlberg et al. (dashed lines) to the AIL population here (solid lines)).

In addition, the approach used here provided sufficient resolution to partition the latter QTL into three independent peaks (4/23, 4/36, 4/70 Chr/Mb, Figure 3, panel B, compare full AIL populations (solid lines) with *deep-Stripes* markers (empty squares, green) to the marker panel used in Wahlberg et al. (full circles, pink) to see how the resolution helped separate 4/23 and 4/36).

#### Large contributions by mapped QTL to the selection response

The AIL was produced by intercrossing chickens from HWS and LWS after 40 generations of selection, and the founders for the intercross differed more than eight-fold (1,341g) in 56-day weights (Table 1). Estimated from the AIL F_2_ - F_18_, the 12 QTL reaching genome wide significance together explain 501.4g (37.4%) of the difference between the parental lines (Table 1), improving upon the 159g (12%) explained by the two significant QTL in Wahlberg et al. (2009). When also considering suggestive QTL (Table S1), the 25 QTL detected here explain 729g (54.4%) of the parental difference compared to the 227.4g (17%) for the 2+1 significant and suggestive QTL in Wahlberg et al. (2009). Together, the significant (suggestive) QTL mapped here/by Wahlberg et al. (2009) explained 8.3/5.2% (11.1/7.1%) of the residual phenotypic variance in the AIL-*F*_2_-*F*_18_ population (Table 1), whereas the 42 peak markers derived from the FDR-approach explain 14.6% of the residual phenotypic variance, or 1,130g, representing 84.3% of the difference between the founding lines.

#### Concordance with previous estimates of effect size

The difference in effect size estimates for *Growth1* on Chromosome 1 between Wahlberg et al. (2009) and the approach used here were within the range of the standard error (34.2 ± 9.2g and 31.6 ± 4.1g, respectively). The effect size estimated here for *Growth6* (20.7 ± 4.4g) on Chromosome 4 was lower than the previous estimate (36.3 ± 8.3g). However, the current approach found a total of 3+1 significant and suggestive QTL on Chromosome 4, of which two significant and one suggestive QTL were within a QTL region (*Growth7*) reported in the first analysis of the F_2_ generation (Jacobsson et al. 2005), but that later showed no significant association with 56-day body weight in the extended analysis of the same individuals with a more comprehensive marker set by Wahlberg et al. (2009). Taken together, these QTL explained a total of 81.2g. In contrast, *Growth9* on Chromosome 7 had a reduced effect size estimate (25.6 ± 4.0g), compared to the 43.2 ± 9g estimated by Wahlberg et al. (2009).

### Additional power in the analyses facilitates detection of new QTL

#### A novel QTL on chromosome 1 revealed by combining data across generations

A genome-wide significant QTL was mapped to 27 Mb on Chromosome 1 (Figure 3a; Table 1). This region was covered by multiple SNP and microsatellite markers in the earlier genome-scans (Wahlberg et al. 2009; Figure 1). However, as these markers segregated in the founder lines, the estimation of founder-line QTL genotypes was less precise than the current approach utilizing low-coverage sequencing data (Figure 3c). To evaluate the potential contribution by either improved marker density or increased number of individuals to an increase in statistical power, the genome scan was performed on different subsets of the data. The first subset was F_2_ individuals from Wahlberg et al. (2009), and second, the subset of bins containing SNP and microsatellite markers from the same study. Thirdly, all combinations with the individuals and markers from this study were used. The results indicated (Figure 3) that the discovery of this QTL was driven by the integration of the AIL and not an increase in marker density.

#### New QTL revealed in regions with poor marker coverage in the genome

The low coverage sequencing approach implemented here improved the genome-wide marker coverage in this population. In particular, the coverage was improved by reaching further out the ends of chromosomes on almost all chromosomes, but specifically 5, 6, and 8 (Figure 1). As a result, a QTL for 56-day body weight was revealed on Chromosome 8 where an additional 7 Mb (0-7 Mb) was covered on its distal end (Figure 4b). The QTL peak is located on the end of the chromosome and does not extend into the part of the chromosome covered in the earlier studies (Wahlberg et al. 2009). Its effect is relatively small, and hence explains a modest amount of the variance in 56-day body weight (Table S1). Its peak location was further located approximately 20 cM outside of the most distal marker in the linkage map used by Wahlberg et al. (2009). Figure 4 illustrates how the combination of better regional coverage and gain in power from merging individuals from multiple generations in the AIL resulted in its detection.

**Figure 4:**
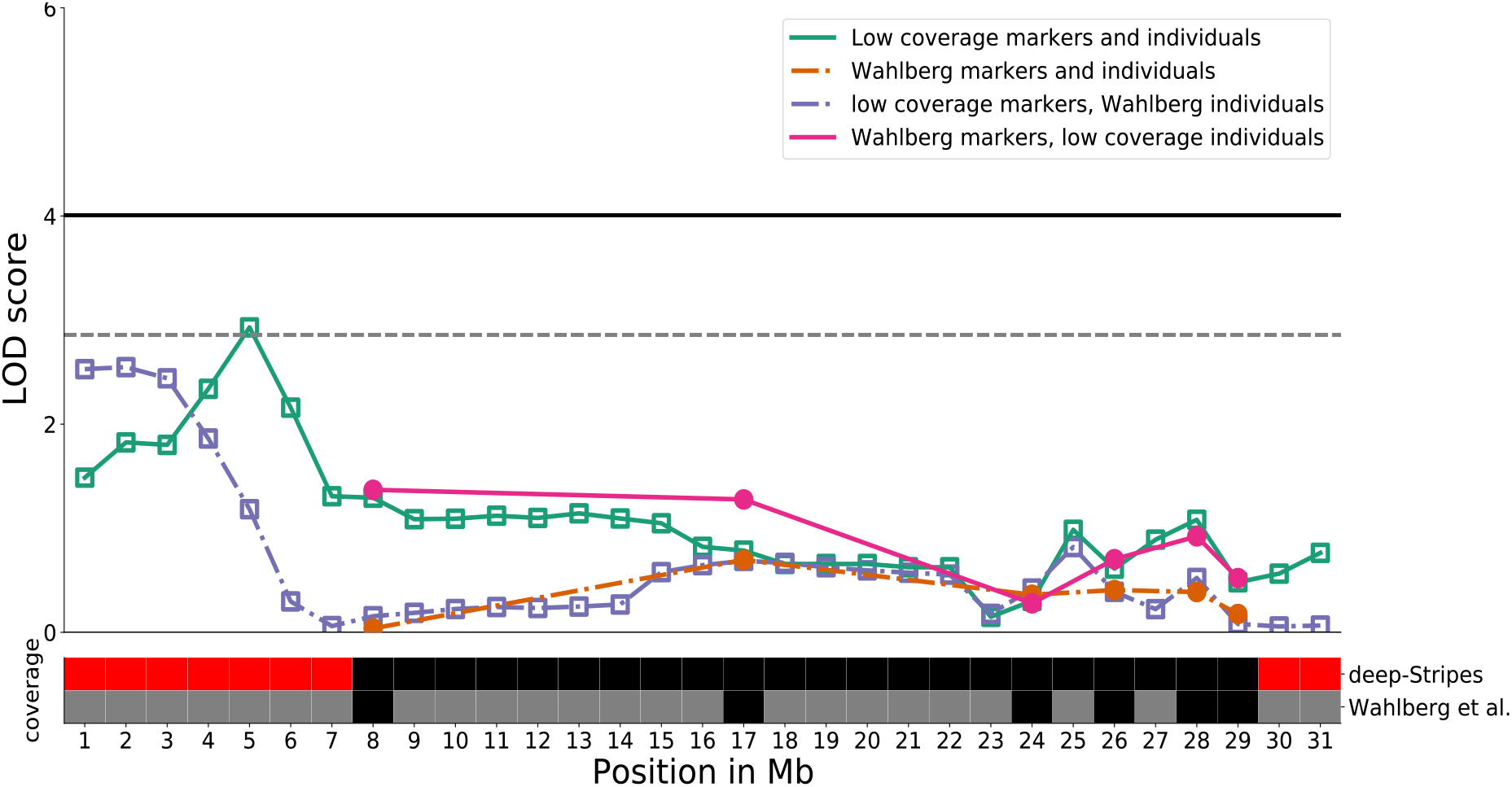
Upper panel: QTL scan for 56-day body weight across Chromosome 8, using the individuals from the original *F*_2_ genome scan (Wahlberg et al. 2009, dashed lines) and all AIL individuals (solid lines) utilising only the marker positions from Wahlberg et al. (filled circles) or all low-coverage markers (empty squares). **Lower panel:** Chromosome 8 sectioned into 1 Mb bins, indicating whether markers are present (black, red) or not present (grey) for both low coverage data (upper row) and the marker set from the original *F*_2_ scan by Wahlberg *et al.* (2009; lower row) Red highlights chromosome ends with new genotypes outside the outermost markers in Wahlberg et al. (2009).

## Discussion

QTL mapping in experimental intercrosses is a valuable strategy for the detection of loci contributing to differences in complex traits between founder populations. Various F_2_ populations have been developed and analysed (e.g. Wright et al. 2006; Kukekova et al. 2011; Solberg Woods 2013; Ying Guo et al. 2016), and in some cases deep intercross populations were bred to increase resolution of the mapped regions. In crosses between segregating, outbred populations, power is often limited by a shortage of between-population informative markers. As a result, low information regions were often relatively poorly explored. Examples include chromosome ends, microchromosomes, and even lowly differentiated intrachromosomal regions. Unfortunately, few populations have been re-genotyped with high density markers due to the associated high costs. Here, we have implemented and evaluated a cost- and time-efficient genotyping strategy utilizing low-coverage sequencing to increase the genome-wide coverage in QTL scans. A software implementation is provided for genotype estimation in F_2_ and deep intercrosses between outbred founder populations. A large advanced intercross chicken line was analysed to empirically illustrate the value of increasing genotype density and quality, as well as making use of available samples across generations, to improve the power in QTL mapping.

### By doubling the number of genome-wide significant QTL identified, three times as much of the founder-line difference for the selected trait is now explained by confidently identified loci

In this study, the intercross population analysed was bred from the Virginia body weight lines. These two pedigreed populations were divergently selected long-term for a single trait, 56-day body weight, for 41 generations before the intercross was formed. Earlier whole-genome analyses of this intercross were challenged to detect genome-wide significant QTL. Only one QTL affecting the selected trait reached genome wide significance in the first analyses that covered ~80% of the genome with 145 markers (Jacobsson et al. 2005). In a subsequent study, utilizing a denser genetic map covering ~93% of the genome with 434 markers (Wahlberg et al. 2009), two genome-wide significant QTL associated with the same trait were found. Fine-mapping analyses in later generations of the intercross focusing first on significant and suggestive QTL regions (Besnier et al. 2011; Pettersson et al. 2011), and later also on selective-sweep regions detected in comparisons between the divergent founder lines (Johansson et al. 2010; Sheng et al. 2015; Pettersson et al. 2013; Zan et al. 2017), all suggested that the genetic architecture of 56-day body weight in this population was highly polygenic and that individual loci contributed small marginal effects.

The increased power in our study obtained by increasing quality and coverage of markers, as well as increasing population-size by integrating across generational data, facilitated detection of nine additional QTL associated with 56-day body weight at a genome wide significance leve, which had not been identified as significant or suggestive QTL by Wahlberg et al. (2009). The residual phenotypic variance explained by the mapped QTL increased from 5.3% by the two genome-wide significant QTL mapped by Wahlberg et al (2009) to 8.3% by the 12 genome-wide significant QTL detected here. The effect sizes of the QTL estimated in the population analysed here were often lower than earlier estimates, with the exception of *Growth1*. The reduced effect size for *Growth9* was not surprising, given that Chromosome 7, and *Growth9* in particular, were previously found to be involved in many epistatic interactions (M. Pettersson et al. 2011) and is likely a complex region. As such, the effect size estimate in the AIL was likely affected by fluctuations in allele-frequency not present in the F_2_ populations. For *Growth6*, while the estimate is lower than that in Wahlberg et al. (2009), when taking into account all significant and suggestive QTL found on Chromosome 4, the sum of the effect sizes was much larger and almost identical (81.2g compared to 79.1g) to the estimate for all regions on Chromosome 4, namely *Growth6* and *Growth7,* made by Jacobsson et al. (2005)., This similarity suggests that our study captures and confirms the same effect on body weight where only suggestive or circumstantial evidence from related phenotypes was previously available. Additionally, this study enabled mapping it more precisely onto multiple independent regions and confirms the existence of multiple regions previously associated with 56-day body weight in the wake of selection-scans (Zan et al. 2017). In total, 3 of the selection markers outside of previously identified QTL regions overlap with the significant or suggestive QTL in this study. Additionally, out of eleven total markers significant in Zan et al. (2017), all but two were correlated with elevated LOD scores. This is consistent with the 20% FDR threshold employed by Zan et al. (2017). Evaluating these regions under a more lenient threshold corroborates this, as most, if not all, of these elevated regions retain significance when accounting for multiple testing leniently. Further, the considerable overlap between these QTL and the previously identified sweep regions from Johansson et al. (2005), not only supports the thesis that these are real, marginal QTL, but also lends credence to previous studies.

Overall, the effects of the significant QTL contributed 37.4% of the founder-line difference (Table S2), which is 3.1 times the founder-line difference explained by the significant QTL in Wahlberg et al. (2009). While power was sufficient to triple the founder-line difference explained, for significant and suggestive QTL taken together, these QTL only explain 54.4% of the founder-line difference, indicating that it likely is still too low to capture the full genetic architecture of the trait studied. Relaxing the association threshold further, the QTL peak markers significant at an 10% FDR-threshold explain more than 84% of the difference between the founder lines. While this may suggest that we are getting closer to identifying the majority of loci contributing to the weight difference, given the large fraction of the genome covered by these markers, the approach used here to estimate the effect size of individual loci likely leads to a slight overestimation from residual linkage between QTL and tagging of linked loci.

Similarly, 30.3% of the bins on the covered autosomal chromosomes are statistically associated with the phenotype after lenient adjustment for multiple testing. This means that any overlap between the extended QTL and the selective sweep regions has to be interpreted with caution. Still, this is equally a testament of the polygenicity of the investigated trait, and it is reasonable to believe that these regions were undergoing true and detectable selection, given the strong single trait artificial selection regime in the selected lines sustained over 40 generations (Siegel 1962).

### Added value by an increased genome-wide marker coverage

Genotyping by sequencing has provided opportunities to both increase marker density and decrease the cost for genotyping. Strategies based on low-coverage sequencing approaches open cost-efficient opportunities to re-analyse existing experimental populations. Here, we use one such approach targeted to deep outbred intercrosses. This approach increased the overall marker coverage of the chicken genome from 93% in the latest study by Wahlberg et al. (2009), as estimated as coverage of the autosomal genetic map, to 1,058 Mb (99.3% of 1,065 Mb, Genome Reference Consortium 2018) with a mean density of 102 line-informative markers per Mb.

Outbred founder lines are beneficial for the density of line-informative markers, because still segregating markers are line-informative due to specific ancestor combinations. However, as the number of these markers shrinks with the increase in unique ancestors contributing to each individual in later generations, the depth of generations one can investigate with this can be limited by the number of founding individuals as well as the size of each generation. Thus, it is important that the founder lines are sufficiently divergent to provide a minimum resolution via fixed markers. However, because the non-physical window size of the imputation process does not sacrifice higher resolution in divergent regions, it is likely that any cross between lines with a significant genetic component to their phenotypic divergence will have sufficient coverage with informative markers in regions of interest. For the AIL used in this study, it is likely that the resolution is not limited by the marker coverage, but rather by both the 1Mb binning approach and the 50/200 marker sliding window imputation of founder genotypes. This is because there were enough accumulated recombinations in the later generations that were lost through these approaches. While these were chosen for robustness and provided an appropriate resolution/power trade-off across all individuals, modifying these parameters could provide added opportunities for fine mapping.

The increased marker density also increased the information content throughout the genome, further increasing power in the QTL analyses. In particular, coverage was extended at the ends of the larger chromosomes. This led to locating a novel suggestive QTL on Chromosome 8 that was primarily due to increase in coverage, though the increase in power helped elevate and define the peak of the QTL. It was seen in the full marker set using only the F_2_ population, but without a definitive peak. This is likely due to a lack of recombination events distal to the peak in the F_2_ population, though the LOD scores for the outermost 3 Mb were above the 5% permutation threshold for chromosome-wide significance.

In addition, two additional small linkage groups were also covered. The minor regions of the genome that remain are present in scaffolds containing too few markers for reliable genotype estimation using the *Stripes* pipeline. Additional work beyond this study is needed to estimate the genotype in these regions using alternative genotyping or bioinformatics approaches before they can be included in QTL mapping studies.

### Integrating across-population data

Integrating data across an AIL population provides value to the QTL scan, by helping to map new QTL and refine the resolution and explanatory power of existing ones. This is particularly so if intermediate generations already exist due to previous attempts in fine mapping. At less than 1 EUR/sample (Zan et al. 2019), the approach demonstrated here provides a cost-effective approach to enhancing the statistical power to dissect complex traits from potentially any experimental population or selection experiment.

For AIL populations with smaller intermediate generations and multiple siblings or half sibs, correcting generation with a mixed or fixed effect model will likely result in overcorrection due to the correlation between generations and kinship. Standardisation using z-scores provides an acceptable trade-off between overcorrecting and confidence in accounting for generational batch effects.

In conclusion, this study represents the most comprehensive study of the individual loci forming the genetic basis of the highly polygenic, long-term selection responses on 56-day body weight in the Virginia chicken lines to date. It contributes not only to our current understanding of the genetic basis of body weight in chickens, but also provides a solid methodological foundation to further investigate the genetic architecture of complex traits in populations with similar design.

## Acknowledgements

We thank Lars Rönnegård for the helpful comments and discussions regarding the stability of the minimisation algorithm. The computations and data handling were enabled by resources in projects SNIC 2017/7-53, SNIC 2018-3-170 and SNIC 2020-5-14 provided by the Swedish National Infrastructure for Computing (SNIC) at UPPMAX, partially funded by the Swedish Research Council through grant agreement no. 2018-05973. The work was supported by the Swedish Research Council (grants 349-2005-8628, 621-2012-4634, 2017-3726 and 2018-5991) and FORMAS (grants 2013-450 and 2017-415).

## References

Altshuler, David, Victor J. Pollara, Chris R. Cowles, William J. Van Etten, Jennifer Baldwin, Lauren Linton, and Eric S. Lander. 2000. “An SNP Map of the Human Genome Generated by Reduced Representation Shotgun Sequencing.” Nature 407 (6803): 513–16. https://doi.org/10.1038/35035083.

Andersson, Leif. 2001. “Genetic Dissection of Phenotypic Diversity in Farm Animals.” Nature Reviews Genetics 2 (2): 130–38. https://doi.org/10.1038/35052563.

Andersson, Leif, and Michel Georges. 2004. “Domestic-Animal Genomics: Deciphering the Genetics of Complex Traits.” Nature Reviews Genetics 5 (3): 202–12. https://doi.org/10.1038/nrg1294.

Andolfatto, Peter, Dan Davison, Deniz Erezyilmaz, Tina T. Hu, Joshua Mast, Tomoko Sunayama-Morita, and David L. Stern. 2011. “Multiplexed Shotgun Genotyping for Rapid and Efficient Genetic Mapping.” Genome Research 21 (4): 610–17. https://doi.org/10.1101/gr.115402.110.

Auwera, Geraldine A. Van der, Mauricio O. Carneiro, Christopher Hartl, Ryan Poplin, Guillermo del Angel, Ami Levy-Moonshine, Tadeusz Jordan, et al. 2013. “From FastQ Data to High-Confidence Variant Calls: The Genome Analysis Toolkit Best Practices Pipeline.” Current Protocols in Bioinformatics 43 (1): 11.10.1-11.10.33. https://doi.org/10.1002/0471250953.bi1110s43.

Benjamini, Yoav, and Yosef Hochberg. 1995. “Controlling the False Discovery Rate: A Practical and Powerful Approach to Multiple Testing.” Journal of the Royal Statistical Society. Series B (Methodological) 57 (1): 289–300.

Besnier, Francois, Per Wahlberg, Lars Rönnegård, Weronica Ek, Leif Andersson, Paul B. Siegel, and Orjan Carlborg. 2011. “Fine Mapping and Replication of QTL in Outbred Chicken Advanced Intercross Lines.” Genetics Selection Evolution 43 (1): 3. https://doi.org/10.1186/1297-9686-43-3.

Boyle, Evan A., Yang I. Li, and Jonathan K. Pritchard. 2017. “An Expanded View of Complex Traits: From Polygenic to Omnigenic.” Cell 169 (7): 1177–86. https://doi.org/10.1016/j.cell.2017.05.038.

Brandt, Monika, Muhammad Ahsan, Christa F Honaker, Paul B Siegel, and Örjan Carlborg. 2017. “Imputation-Based Fine-Mapping Suggests That Most QTL in an Outbred Chicken Advanced Intercross Body Weight Line Are Due to Multiple, Linked Loci.” G3 Genes|Genomes|Genetics 7 (1): 119–28. https://doi.org/10.1534/g3.116.036012.

Broman, Karl W., Hao Wu, Saunak Sen, and Gary A. Churchill. 2003. “R/Qtl: QTL Mapping in Experimental Crosses.” Bioinformatics (Oxford, England) 19 (7): 889–90. https://doi.org/10.1093/bioinformatics/btg112.

Carlborg, Örjan, Lina Jacobsson, Per Åhgren, Paul Siegel, and Leif Andersson. 2006. “Epistasis and the Release of Genetic Variation during Long-Term Selection.” Nature Genetics 38 (4): 418–20. https://doi.org/10.1038/ng1761.

Cock, Peter J. A., Tiago Antao, Jeffrey T. Chang, Brad A. Chapman, Cymon J. Cox, Andrew Dalke, Iddo Friedberg, et al. 2009. “Biopython: Freely Available Python Tools for Computational Molecular Biology and Bioinformatics.” Bioinformatics 25 (11): 1422–23. https://doi.org/10.1093/bioinformatics/btp163.

DePristo, Mark A., Eric Banks, Ryan Poplin, Kiran V. Garimella, Jared R. Maguire, Christopher Hartl, Anthony A. Philippakis, et al. 2011. “A Framework for Variation Discovery and Genotyping Using Next-Generation DNA Sequencing Data.” Nature Genetics 43 (5): 491–98. https://doi.org/10.1038/ng.806.

Dudley, J. W. 2007. “From Means to QTL: The Illinois Long-Term Selection Experiment as a Case Study in Quantitative Genetics.” Crop Science 47 (S3): S-20–S-31. https://doi.org/10.2135/cropsci2007.04.0003IPBS.

Flint, Jonathan, and Richard Mott. 2001. “Finding the Molecular Basis of Quatitative Traits: Successes and Pitfalls.” Nature Reviews Genetics 2 (6): 437–45. https://doi.org/10.1038/35076585.

Forsberg, Simon K. G., Joshua S. Bloom, Meru J. Sadhu, Leonid Kruglyak, and Örjan Carlborg. 2017. “Accounting for Genetic Interactions Improves Modeling of Individual Quantitative Trait Phenotypes in Yeast.” Nature Genetics 49 (4): 497–503. https://doi.org/10.1038/ng.3800.

Genome Reference Consortium. 2018. “Chicken Genome - Assembly GRC6a.” 2018. https://www.ncbi.nlm.nih.gov/assembly/GCF_000002315.5/.

Groenen, Martien A. M., Per Wahlberg, Mario Foglio, Hans H. Cheng, Hendrik-Jan Megens, Richard P. M. A. Crooijmans, Francois Besnier, et al. 2009. “A High-Density SNP-Based Linkage Map of the Chicken Genome Reveals Sequence Features Correlated with Recombination Rate.” Genome Research 19 (3): 510–19. https://doi.org/10.1101/gr.086538.108.

Groenen, Martien A.M., Hans H. Cheng, Nat Bumstead, Bernard F. Benkel, W. Elwood Briles, Terry Burke, Dave W. Burt, et al. 2000. “A Consensus Linkage Map of the Chicken Genome.” Genome Research 10 (1): 137–47.

Guo, Ying, Mette Lillie, Yanjun Zan, Jeanette Beranger, Alison Martin, Christa. F. Honaker, Paul. B. Siegel, and Örjan Carlborg. 2019. “A Genomic Inference of the White Plymouth Rock Genealogy.” Poultry Science 98 (11): 5272–80. https://doi.org/10.3382/ps/pez411.

Guo, Ying, Xiaorong Gu, Zheya Sheng, Yanqiang Wang, Chenglong Luo, Ranran Liu, Hao Qu, et al. 2016. “A Complex Structural Variation on Chromosome 27 Leads to the Ectopic Expression of HOXB8 and the Muffs and Beard Phenotype in Chickens.” PLOS Genetics 12 (6): e1006071. https://doi.org/10.1371/journal.pgen.1006071.

Hopkins, Cyril. 1899. “Improvement in the Chemical Composition of the Corn Kernel.” Journal of the American Chemical Society. https://pubs.acs.org/doi/pdf/10.1021/ja02061a012?casa_token=HKkURz-FUlwAAAAA%3AS1Zyocbjg7xnN7U_gv4E4qXmvqfL1w5pTHlXcDFNgKVs6OGsPO-4JMd2PnAIRkqwwRlkmvRhL2E4Oxs&.

Jacobsson, Lina, Hee-Bok Park, Per Wahlberg, Robert Fredriksson, Miguel Perez-Enciso, Paul B. Siegel, and Leif Andersson. 2005. “Many QTLs with Minor Additive Effects Are Associated with a Large Difference in Growth between Two Selection Lines in Chickens.” Genetics Research 86 (2): 115–25. https://doi.org/10.1017/S0016672305007767.

Johansson, Anna M., Mats E. Pettersson, Paul B. Siegel, and Örjan Carlborg. 2010. “Genome-Wide Effects of Long-Term Divergent Selection.” PLOS Genetics 6 (11): e1001188. https://doi.org/10.1371/journal.pgen.1001188.

Knott, Sara A., and Chris S. Haley. 1992. “Maximum Likelihood Mapping of Quantitative Trait Loci Using Full-Sib Families.” Genetics 132 (4): 1211–22.

Knott, Sara A., Lena Marklund, Chris S. Haley, Kjell Andersson, William Davies, Hans Ellegren, Merete Fredholm, et al. 1998. “Multiple Marker Mapping of Quantitative Trait Loci in a Cross between Outbred Wild Boar and Large White Pigs.” Genetics 149 (2): 1069–80.

Kukekova, Anna V., Lyudmila N. Trut, Kevin Chase, Anastasiya V. Kharlamova, Jennifer L. Johnson, Svetlana V. Temnykh, Irina N. Oskina, et al. 2011. “Mapping Loci for Fox Domestication: Deconstruction/Reconstruction of a Behavioral Phenotype.” Behavior Genetics 41 (4): 593–606. https://doi.org/10.1007/s10519-010-9418-1.

Lango Allen, Hana, Karol Estrada, Guillaume Lettre, Sonja I. Berndt, Michael N. Weedon, Fernando Rivadeneira, Cristen J. Willer, et al. 2010. “Hundreds of Variants Clustered in Genomic Loci and Biological Pathways Affect Human Height.” Nature 467 (7317): 832–38. https://doi.org/10.1038/nature09410.

Li, Heng. 2011. “A Statistical Framework for SNP Calling, Mutation Discovery, Association Mapping and Population Genetical Parameter Estimation from Sequencing Data.” Bioinformatics (Oxford, England) 27 (21): 2987–93. https://doi.org/10.1093/bioinformatics/btr509.

Li, Heng. 2013. “Aligning Sequence Reads, Clone Sequences and Assembly Contigs with BWA-MEM.” ArXiv:1303.3997 [q-Bio], May. http://arxiv.org/abs/1303.3997.

Li, Heng, Bob Handsaker, Alec Wysoker, Tim Fennell, Jue Ruan, Nils Homer, Gabor Marth, Goncalo Abecasis, Richard Durbin, and 1000 Genome Project Data Processing Subgroup. 2009. “The Sequence Alignment/Map Format and SAMtools.” Bioinformatics (Oxford, England) 25 (16): 2078–79. https://doi.org/10.1093/bioinformatics/btp352.

Lillie, M, Z Y Sheng, C F Honaker, L Andersson, P B Siegel, and Ö Carlborg. 2018. “Genomic Signatures of 60 Years of Bidirectional Selection for 8-Week Body Weight in Chickens.” Poultry Science 97 (3): 781–90. https://doi.org/10.3382/ps/pex383.

Lillie, Mette, Christa F Honaker, Paul B Siegel, and Örjan Carlborg. 2019. “Bidirectional Selection for Body Weight on Standing Genetic Variation in a Chicken Model.” G3 Genes|Genomes|Genetics 9 (4): 1165–73. https://doi.org/10.1534/g3.119.400038.

Mackay, Trudy F. C. 2001. “The Genetic Architecture of Quantitative Traits.” Annual Review of Genetics 35 (1): 303–39. https://doi.org/10.1146/annurev.genet.35.102401.090633.

Mackay, Trudy F. C. 2014. “Epistasis and Quantitative Traits: Using Model Organisms to Study Gene–Gene Interactions.” Nature Reviews Genetics 15 (1): 22–33. https://doi.org/10.1038/nrg3627.

McKenna, Aaron, Matthew Hanna, Eric Banks, Andrey Sivachenko, Kristian Cibulskis, Andrew Kernytsky, Kiran Garimella, et al. 2010. “The Genome Analysis Toolkit: A MapReduce Framework for Analyzing next-Generation DNA Sequencing Data.” Genome Research 20 (9): 1297–1303. https://doi.org/10.1101/gr.107524.110.

Pedersen, Brent S., and Aaron R. Quinlan. 2017. “Cyvcf2: Fast, Flexible Variant Analysis with Python.” Bioinformatics 33 (12): 1867–69. https://doi.org/10.1093/bioinformatics/btx057.

Peirce, Jeremy L., Hongqiang Li, Jintao Wang, Kenneth F. Manly, Robert J. Hitzemann, John K. Belknap, Glenn D. Rosen, et al. 2006. “How Replicable Are MRNA Expression QTL?” Mammalian Genome 17 (6): 643–56. https://doi.org/10.1007/s00335-005-0187-8.

Pértille, Fábio, Carlos Guerrero-Bosagna, Vinicius Henrique da Silva, Clarissa Boschiero, José de Ribamar da Silva Nunes, Mônica Corrêa Ledur, Per Jensen, and Luiz Lehmann Coutinho. 2016. “High-Throughput and Cost-Effective Chicken Genotyping Using Next-Generation Sequencing.” Scientific Reports 6 (1): 26929. https://doi.org/10.1038/srep26929.

Pettersson, Mats, Francois Besnier, Paul B. Siegel, and Örjan Carlborg. 2011. “Replication and Explorations of High-Order Epistasis Using a Large Advanced Intercross Line Pedigree.” PLOS Genetics 7 (7): e1002180. https://doi.org/10.1371/journal.pgen.1002180.

Pettersson, Mats E., Anna M. Johansson, Paul B. Siegel, and Örjan Carlborg. 2013. “Dynamics of Adaptive Alleles in Divergently Selected Body Weight Lines of Chickens.” G3: Genes, Genomes, Genetics 3 (12): 2305–12. https://doi.org/10.1534/g3.113.008375.

R Core Team, R Core Team. 2013. “R: A Language and Environment for Statistical Computing.”

Rowan, Beth A., Vipul Patel, Detlef Weigel, and Korbinian Schneeberger. 2015. “Rapid and Inexpensive Whole-Genome Genotyping-by-Sequencing for Crossover Localization and Fine-Scale Genetic Mapping.” G3: Genes, Genomes, Genetics 5 (3): 385–98. https://doi.org/10.1534/g3.114.016501.

Seabold, Skipper, and Josef Perktold. 2010. “Statsmodels: Econometric and Statistical Modeling with Python.” In, 92–96. Austin, Texas. https://doi.org/10.25080/Majora-92bf1922-011.

Sheng, Zheya, Mats E. Pettersson, Christa F. Honaker, Paul B. Siegel, and Örjan Carlborg. 2015. “Standing Genetic Variation as a Major Contributor to Adaptation in the Virginia Chicken Lines Selection Experiment.” Genome Biology 16 (1): 219. https://doi.org/10.1186/s13059-015-0785-z.

Siegel, P. B. 1962. “Selection for Body Weight at Eight Weeks of Age: 1. Short Term Response and Heritabilities1.” Poultry Science 41 (3): 954–62. https://doi.org/10.3382/ps.0410954.

Solberg Woods, Leah C. 2013. “QTL Mapping in Outbred Populations: Successes and Challenges.” Physiological Genomics 46 (3): 81–90. https://doi.org/10.1152/physiolgenomics.00127.2013.

Wahlberg, Per, Örjan Carlborg, Mario Foglio, Xavier Tordoir, Ann-Christine Syvänen, Mark Lathrop, Ivo G. Gut, Paul B. Siegel, and Leif Andersson. 2009. “Genetic Analysis of an F2 Intercross between Two Chicken Lines Divergently Selected for Body-Weight.” BMC Genomics 10 (1): 248. https://doi.org/10.1186/1471-2164-10-248.

Whalen, Andrew, Roger Ros-Freixedes, David L. Wilson, Gregor Gorjanc, and John M. Hickey. 2018. “Hybrid Peeling for Fast and Accurate Calling, Phasing, and Imputation with Sequence Data of Any Coverage in Pedigrees.” Genetics Selection Evolution 50 (1): 67. https://doi.org/10.1186/s12711-018-0438-2.

Wood, Andrew R., Tonu Esko, Jian Yang, Sailaja Vedantam, Tune H. Pers, Stefan Gustafsson, Audrey Y. Chu, et al. 2014. “Defining the Role of Common Variation in the Genomic and Biological Architecture of Adult Human Height.” Nature Genetics 46 (11): 1173–86. https://doi.org/10.1038/ng.3097.

Wright, D., S. Kerje, K. Lundström, J. Babol, K. Schütz, P. Jensen, and L. Andersson. 2006. “Quantitative Trait Loci Analysis of Egg and Meat Production Traits in a Red Junglefowl × White Leghorn Cross.” Animal Genetics 37 (6): 529–34. https://doi.org/10.1111/j.1365-2052.2006.01515.x.

Yang, Jian, Beben Benyamin, Brian P. McEvoy, Scott Gordon, Anjali K. Henders, Dale R. Nyholt, Pamela A. Madden, et al. 2010. “Common SNPs Explain a Large Proportion of the Heritability for Human Height.” Nature Genetics 42 (7): 565–69. https://doi.org/10.1038/ng.608.

Zan, Yanjun, and Örjan Carlborg. 2020. “Dynamic Genetic Architecture of Yeast Response to Environmental Perturbation Shed Light on Origin of Cryptic Genetic Variation.” PLOS Genetics 16 (5): e1008801. https://doi.org/10.1371/journal.pgen.1008801.

Zan, Yanjun, Thibaut Payen, Mette Lillie, Christa F. Honaker, Paul B. Siegel, and Örjan Carlborg. 2019. “Genotyping by Low-Coverage Whole-Genome Sequencing in Intercross Pedigrees from Outbred Founders: A Cost-Efficient Approach.” Genetics Selection Evolution 51 (1): 44. https://doi.org/10.1186/s12711-019-0487-1.

Zan, Yanjun, Zheya Sheng, Mette Lillie, Lars Rönnegård, Christa F. Honaker, Paul B. Siegel, and Örjan Carlborg. 2017. “Artificial Selection Response Due to Polygenic Adaptation from a Multilocus, Multiallelic Genetic Architecture.” Molecular Biology and Evolution 34 (10): 2678–89. https://doi.org/10.1093/molbev/msx194.

Zhang, X., P. Pérez-Rodríguez, K. Semagn, Y. Beyene, R. Babu, M. A. López-Cruz, F. San Vicente, et al. 2015. “Genomic Prediction in Biparental Tropical Maize Populations in Water-Stressed and Well-Watered Environments Using Low-Density and GBS SNPs.” Heredity 114 (3): 291–99. https://doi.org/10.1038/hdy.2014.99.

