## supplementary tables and figures for "Low-coverage sequencing in a deep intercross of the Virginia body weight lines provides insight to the polygenic genetic architecture of growth: novel loci revealed by increased power and improved genome-coverage"

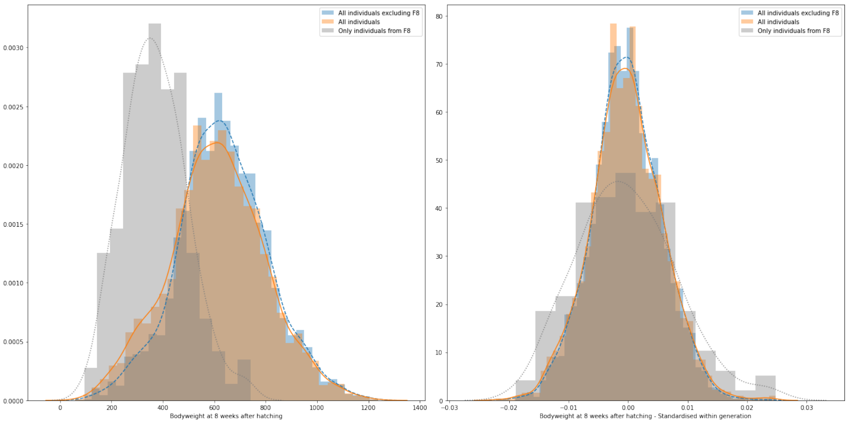

**Figure S1:** Standardisation of 56-day body weight. Left panel shows body weight before standardisation, right panel shows standardised 56-day body weight. All $F_8$ individuals (Grey), all individuals (Orange), and all individuals excluding $F_8$ (Blue)

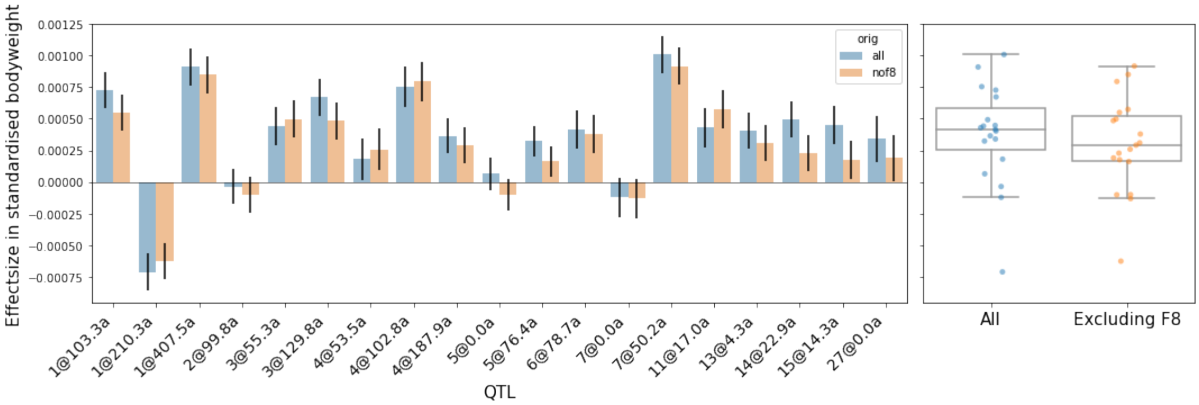

**Figure S2:** Effect Size in standardised body weight, including $F_8$ (blue) and excluding $F_8$ (orange). Left panel shows a barchart of the additive effect at each locus, right panel shows a box/stripplot for all loci combined

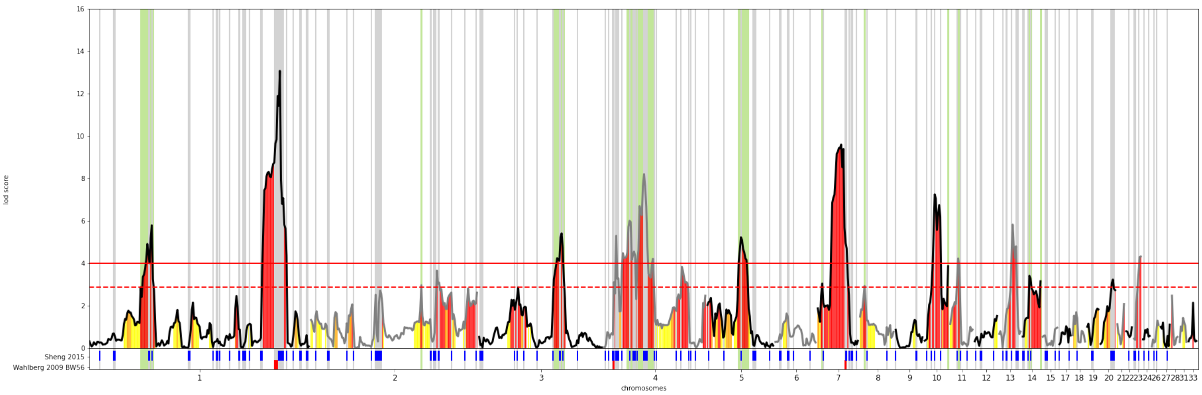

**Figure S3**: Overview of all LOD scores for all investigated chromosomes. Red, orange, and yellow colored vertical sections represent regions retaining significant association to 56-day body weight after adjusting for multiple testing with the benjamini hochberg procedure at an alpha of 1,5, and 10%, respectively. Blue vertical sections below the x-axis are markers indicating selective sweeps from Sheng et al. (2017)

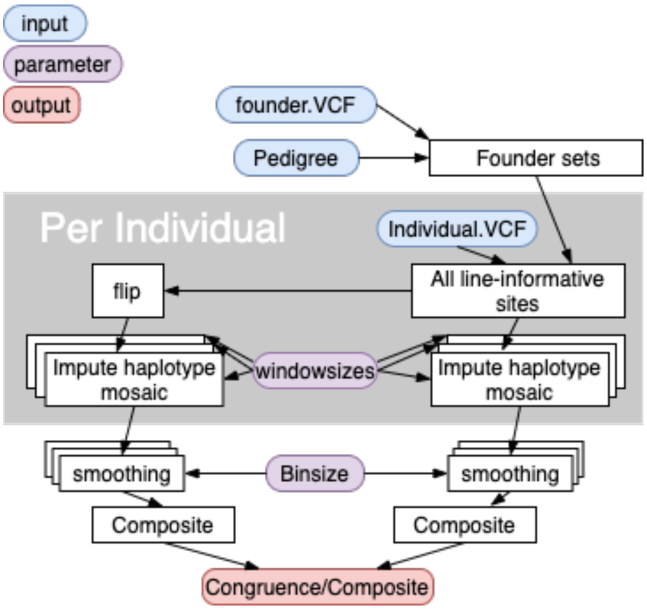

**Figure S4:** Overview of the modified imputation process

**Table S1.** All suggestive and significant QTL peaks for 56-day body weight and overlaps with earlier reported QTL in this population

| | Chromosome | Position(Mb) | a(SE) | %var | lod | QTL | previous |
|---|---|---|---|---|---|---|---|

| | | | | | | | |
|---|---|---|---|---|---|---|---|
| | 1 | 56 | 19.6(4.0) | 0.43 | **5.79** | | |
| | 1 | 171 | 31.6(4.1) | 1.21 | **13.08** | Growth1 | Zan 2017, Wahlberg 2009, Jacobsson 2005 |
| | 2 | 99 | 13.5(4.0) | 0.25 | 2.96 | Growth3 | Jacobsson 2005 |
| | 2 | 113 | 16.6(4.0) | 0.14 | 3.65 | Growth3 | Zan 2017, Jacobsson 2005 |
| | 3 | 35 | 11.6(4.3) | 0.27 | 2.87 | Growth4 | Zan 2017, Jacobsson 2005 |
| | 3 | 74 | 20.3(4.2) | 0.31 | **5.41** | Growth4 | Jacobsson 2005 |
| | 4 | 11 | 20.7(4.4) | 0.28 | **5.29** | Growth6 | Zan 2017, Wahlberg 2009 |
| | 4 | 23 | 20.4(4.0) | 0.07 | **6** | Growth7 ? | Jacobsson 2005 |
| | 4 | 36 | 25.6(4.2) | 0.45 | **8.2** | Growth7 ? | Jacobsson 2005 |
| | 4 | 70 | 14.6(3.9) | 0.2 | 3.83 | Growth7 ? | Zan 2017, Jacobsson 2005 |
| | Z | 1 | 12.9(3.5) | 0.11 | 2.95 | | |
| | Z | 10 | 12.8(3.8) | 0.03 | 3.22 | | |
| | Z | 34 | 14.0(3.4) | 0.05 | 3.8 | | |
| | 5 | 30 | 20.9(4.3) | 0.31 | **5.22** | Growth8 | Jacobsson 2005 |
| | 7 | 4 | 5.2(4.3) | 0.28 | 3.04 | | |
| | 7 | 21 | 25.6(4.0) | 1.26 | **9.6** | Growth9 | Zan 2017, Wahlberg 2009, Jacobsson 2005 |
| | 8 | 4 | -7.1(4.2) | 0.3 | 2.93 | | |
| | 10 | 9 | 22.3(3.9) | 0.27 | **7.25** | | Zan 2017 |
| | 10 | 21 | 15.8(4.3) | 0.1 | 3.86 | | |
| | 11 | 7 | 13.2(4.0) | 0.32 | **4.22** | | |

| | | | | | | | |
|---|---|---|---|---|---|---|---|
| | 13 | 12 | 19.6(3.9) | 0.32 | **5.83** | Growth10 | Jacobsson 2005 |
| | 14 | 6 | 14.0(4.1) | 0.16 | 3.41 | | |
| | 14 | 16 | 16.0(4.3) | 0.01 | 3.15 | | |
| | 20 | 11 | 16.3(4.3) | 0.5 | 3.23 | Growth12 | Zan 2017, Wahlberg 2009 |
| | 23 | 6 | 10.9(4.3) | 0.47 | **4.32** | | Zan 2017 |
| | | | a=407 | | | | |
| | | | 2a=814 | | | | |

**Table S2:** Mean bodyweight and number of individuals per generation

| generation | individuals | BW8 - mean(stdev)g |
|---|---|---|
| HWS-$F_0$ | 29 | 1522(36) |
| LWS-$F_0$ | 30 | 181(5) |
| HWS-LWS difference | | 1341 |
| $F_2$ | 930 | 626(186) |
| $F_3$ | 429 | 696(170) |
| $F_4$ | 112 | 593(131) |
| $F_5$ | 118 | 656(155) |
| $F_6$ | 88 | 770(168) |
| $F_7$ | 41 | 661(178) |
| $F_8$ | 292 | 375(122) |
| $F_9$ | 50 | 710(186) |
| $F_{10}$ | 60 | 674(175) |
| $F_{11}$ | 87 | 579(165) |
| $F_{12}$ | 76 | 603(180) |
| $F_{14}$ | 32 | 730(192) |

| | | |
|---|---|---|
| $F_{15}$ | 818 | 613(162) |
| $F_{16}$ | 89 | 650(174) |
| $F_{17}$ | 45 | 576(165) |
| $F_{18}$ | 87 | 528(148) |
| all | 3354 | 614(188) |

**Table S3:** All markers reaching significance after false-discovery-rate adjustment with alpha=10%

| | Chromosome | Position(Mb) | a(SE) in gram | %var | lod |
|---|---|---|---|---|---|
| | 1 | 35 | 10.32(4.156) | 0.11947 | 1.765 |
| | 1 | 56 | 19.62(4.038) | 0.4807 | 5.788 |
| | 1 | 79 | 2.36(4.191) | 0.20871 | 1.799 |
| | 1 | 93 | -12.34(4.009) | 0.19455 | 2.144 |
| | 1 | 132 | 11.27(3.875) | 0.1573 | 2.457 |
| | 1 | 171 | 31.59(4.059) | 1.11277 | 13.077 |
| | 1 | 185 | -3.88(4.077) | 0.08036 | 1.731 |
| | 2 | 5 | 10.55(3.966) | 0.117 | 1.732 |
| | 2 | 22 | -10.7(4.064) | 0.17337 | 1.695 |
| | 2 | 37 | 4.3(4.016) | 0.26358 | 2.06 |
| | 2 | 58 | 10.95(4.229) | 0.09418 | 2.513 |
| | 2 | 62 | 11.87(4.118) | 0.01154 | 2.707 |

| | | | | | |
|---|---|---|---|---|---|
| | 2 | 99 | 13.47(3.97) | 0.1638 | 2.963 |
| | 2 | 113 | 16.57(4.039) | 0.12504 | 3.651 |
| | 2 | 141 | 14.19(3.948) | 0.12695 | 2.808 |
| | 3 | 35 | 11.63(4.268) | 0.2559 | 2.867 |
| | 3 | 74 | 20.35(4.165) | 0.2721 | 5.408 |
| | 4 | 11 | 20.71(4.367) | 0.16289 | 5.294 |
| | 4 | 23 | 20.44(3.989) | 0.05897 | 6.003 |
| | 4 | 37 | 24.82(4.213) | 0.63225 | 7.631 |
| | 4 | 71 | 14.39(3.932) | 0.25961 | 3.623 |
| | 4 | 91 | 11.35(4.241) | 0.2026 | 2.468 |
| | 5 | 3 | 5.73(4.055) | 0.29812 | 2.366 |
| | 5 | 11 | 12.24(3.962) | 0.15097 | 2.1 |
| | 5 | 17 | 9.34(4.222) | 0.09418 | 1.82 |
| | 5 | 30 | 20.89(4.27) | 0.22515 | 5.225 |
| | 6 | 10 | 10.95(4.029) | 0.08556 | 1.629 |
| | 7 | 4 | 5.22(4.256) | 0.30981 | 3.042 |
| | 7 | 21 | 25.56(3.98) | 0.97001 | 9.599 |
| | 8 | 4 | -7.1(4.232) | 0.25772 | 2.93 |
| | 10 | 9 | 22.3(3.912) | 0.29332 | 7.249 |
| | 11 | 7 | 13.17(4.036) | 0.2799 | 4.223 |
| | 13 | 12 | 19.62(3.92) | 0.25145 | 5.826 |
| | 14 | 6 | 14.04(4.116) | 0.14015 | 3.408 |
| | 14 | 16 | 16.0(4.274) | 0.02953 | 3.149 |
| | 18 | 3 | -9.74(4.036) | 0.22344 | 1.689 |
| | 19 | 9 | 8.04(4.08) | 0.12524 | 1.858 |

| | | | | | |
|---|---|---|---|---|---|
| | 20 | 11 | 16.34(4.27) | 0.60823 | 3.231 |
| | 21 | 6 | 9.59(3.496) | 0.26196 | 2.075 |
| | 23 | 6 | 10.86(4.284) | 0.35913 | 4.318 |
| | 28 | 0 | 7.9(4.252) | 0.20064 | 2.47 |
| | 33 | 4 | -12.35(4.411) | 0.25283 | 2.138 |
| | | | a = 565.44 | | |
| | | | 2a = 1130.87 | | |
